# Dopamine pathway characterization during the reproductive mode switch in the pea aphid

**DOI:** 10.1101/2020.03.10.984989

**Authors:** Gaël Le Trionnaire, Sylvie Hudaverdian, Gautier Richard, Sylvie Tanguy, Florence Gleonnec, Nathalie Prunier-Leterme, Jean-Pierre Gauthier, Denis Tagu

**Affiliations:** IGEPP, INRAE, Institut Agro, Univ Rennes, 35653, Le Rheu, France

**Keywords:** *Acyrthosiphon pisum*, photoperiodic response, dopamine pathway, spatio-temporal expression, CRISPR-Cas9

## Abstract

Aphids are major pests of most of the crops worldwide. Such a success is largely explained by the remarkable plasticity of their reproductive mode. They reproduce efficiently by viviparous parthenogenesis during spring and summer generating important damage on crops. At the end of the summer, viviparous parthenogenetic females perceive the photoperiod shortening and transduce this signal to their embryos that change their reproductive fate to produce sexual individuals: oviparous females and males. After mating, those females lay cold-resistant eggs. Earlier studies showed that some transcripts coding for key components of dopamine pathway were regulated between long days and short days conditions suggesting that dopamine might be involved in the transduction of seasonal cues prior to reproductive mode switch. In this study, we aimed at going deeper into the characterization of the expression dynamics of this pathway but also in the analysis of its functional role in this context in the pea aphid *Acyrthosiphon pisum*. We first analysed the level of expression of ten genes of this pathway in embryos and larval heads of aphids reared under long days (asexual producers) or short days (sexual producers) conditions. We then performed in situ hybridization experiments to localize in embryos the *ddc* and *pale* transcripts that are coding for two key enzymes in dopamine synthesis. Finally, Using CRISPR-Cas9 mutagenesis in eggs produced after the mating of sexual individuals, we targeted the *ddc* gene. We could observe strong melanization defaults in *ddc* mutated eggs, which confidently mimicked the Drosophila *ddc* phenotype. Nevertheless, such a lethal phenotype did not allow us to validate the involvement of dopamine as a signaling pathway necessary to trigger the reproductive mode switch in embryos.

## Introduction

Aphids are hemipteran insects that adapt their reproductive mode to seasonal variations. They are capable of alternating between asexual and sexual reproduction during their annual life cycle. In spring and summer, they reproduce by viviparous parthenogenesis: each asexual female can produce nearly a hundred of clonal progeny. When autumn starts, asexual females perceive the shortening of the photoperiod, which triggers a switch of the reproductive fate of their embryos ending up with the production of clonal oviparous sexual females and males (which lack one X chromosome compared to females, thus not truly clonal) in their offspring. After mating with males, oviparous sexual females lay fertilized eggs that enter an obligate 3-month diapause over winter before hatching in early spring. The individuals hatching from those eggs are new genetic combinations with the viviparous parthenogenetic female phenotype. Thus, it is at least four different clonal morphs that are produced within the annual life cycle: males, oviparous sexual females, parthenogenetic viviparous females producing a progeny of parthenogenetic viviparous females (under long days, dubbed “virginoparae”), and parthenogenetic viviparous females producing a progeny of males and oviparous sexual females (under short days, dubbed “sexuparae”).

This peculiar adaptation to seasonality for poikilothermous animals involves several trans-generational complex physiological and molecular mechanisms: perception and integration of the photoperiod signal, its neuro-endocrine transduction to the ovaries and orientation of developmental programs for embryos (reviewed in Le Trionnaire et al., 2013 and Ogawa and Miura, 2014). Development of embryos in viviparous parthenogenetic aphid ovaries is continuous: each ovariole contains several developing embryos at different stages of development. Viviparity in aphids thus implies a telescoping of generations: grandmothers contain daughter embryos that already enclose (at least for the most developed) the germ cells of the granddaughters. Therefore, sensing and integration of the photoperiod by the adult females needs a continuous transduction of the signal towards the embryos, across three generations. Nevertheless, it has been strongly suggested that embryos can directly perceive the changes in photoperiod (Lees, 1964 and Le Trionnaire et al., 2009), which reduces (at least under controlled conditions) to two generations the transduction effect, from the sexuparae mother (starting early at embryonic stages) to its offspring.

For several years, our and other groups have been involved in deciphering these mechanisms (see Le Trionnaire et al., 2013 for a review). Transcriptomic analyses allowed the identification of neuro-endocrine pathways that differ in expression between aphids reared under short or long photoperiod. Most of the analyses were performed on the pea aphid (*Acyrthosiphon pisum*) which was the first aphid species to present a large set of genomic resources (IAGC, 2010 and Legeai et al., 2010). One of the unexpected observations derived from those transcriptomic analyses was a wide regulation of the expression of cuticular protein genes: several cuticular protein mRNAs were down regulated in the heads of pea aphid sexuparae females under short days (Le Trionnaire et al., 2009 and Cortes et al., 2008). Most of these cuticular proteins belong to the RR2 family, known to be involved (at least in *Tribolium castaneum* and other insects, Arakane et al., 2009) in the sclerotization of the cuticle, a process that contributes to cuticle structure stabilization (Andersen, 2010); they usually accumulate in the inner part of the cuticle. The cuticle of the pea aphid has been described as made of three layers: the outer epicuticule, the inner epicuticule and the procuticle (Brey et al., 1985). Despite several trials, we did not succeed in localizing regulated-cuticular proteins in the cuticle and we did not notice any clear modification of the structure or width of head cuticules in sexuparae (unpublished observation). Sclerotization involves different molecules, including dopamine. We previously demonstrated that two genes involved in dopamine synthesis (*pale* and *ddc*) were also down regulated in heads of sexuparae under short days, as were the cuticular proteins (Gallot et al., 2010).

At that stage, we made several hypotheses on the putative roles of dopamine in the regulation of different downstream pathways such as i) synaptic function, ii) sclerotization of the cuticle, and iii) melanization of the cuticle (Figure 1), involving for each pathway various proteins and genes. Dopamine is formed after two enzymatic reactions that first convert L-tyrosine into L-DOPA (by a tyrosine hydroxylase encoded by the *pale* gene in *Drosophila melanogaster*) and second, convert L-DOPA into dopamine (by a DOPA decarboxylase encoded by the *ddc* gene in *D. melanogaster*). Dopamine can be transported in synaptic vesicles by different proteins such as the vesicular monoamine transporters Vmat, the vesicle amine transporter vat1 or the monoamine transmembrane transporter prt. For cuticle sclerotization, two acyldopamines (N-β-alanine-dopamine (NBAD) and N-acetyldopamine (NADA)) are formed and incorporated into the cuticular matrix (Andersen, 2010). This involves several enzymes such as an arylalkylamine N-acetyltransferase (encoded by the *aaNAT* gene - also known as Dat in *D. melanogaster*) that converts dopamine into NADA, an aspartate 1-decarboxylase (the *black* gene in *D. melanogaster*) that converts aspartate into β-alanine and the ebony enzyme that finally links β-alanine with dopamine to form NBAD. The consequence is the formation of a hard layer of cuticle often at the outer part of the exoskeleton, covering internal softer cuticle layers. Dopamine can also be oxidized by laccases (multicopper oxidases) and/or phenoloxidases (PO) to form the dopamine melanin involved in cuticle pigmentation, in conjunction in some instances with structural proteins such as the Yellow protein (Arakane et al., 2010).

**Figure 1.**
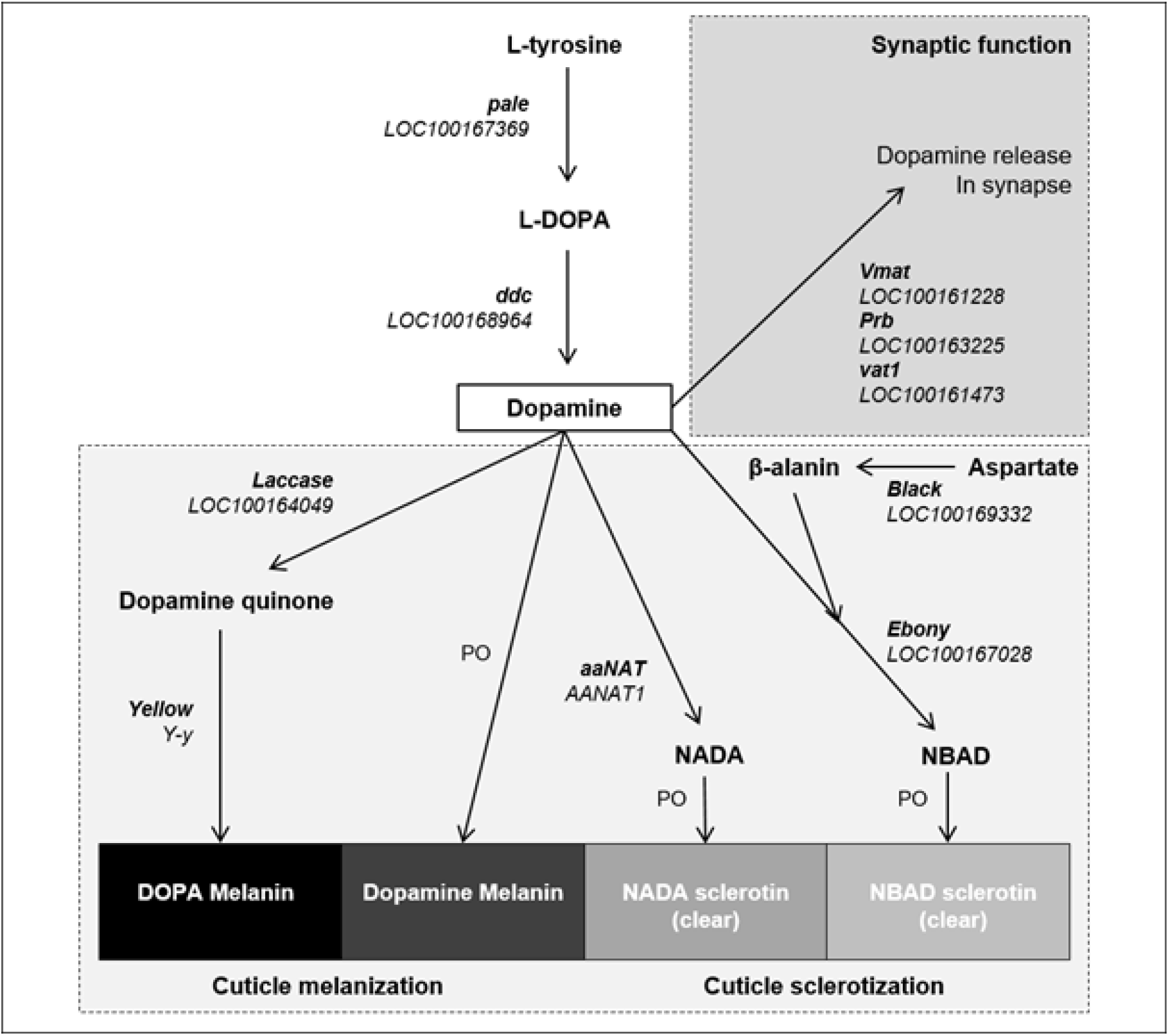
Dopamine biosynthesis and mode of action pathway. L-tyrosine is first hydroxylised by pale (tyrosine hydroxylase) into L-DOPA which is then subsequently decarboxylised by ddc (dopa-decarboxylase) to produce dopamine. Dopamine can then directly be released within synapses and act as a neurotransmitter. Vesicle amine transporters such as Vmat, prb and vat1 are likely to play a key role in this process. Dopamine can also be used as a structural component of the cuticle: either as DOPA melanin (black cuticle) when processed by laccase and yellow enzymes, or as NADA sclerotin (clear cuticle) through the action of aaNAT enzyme, or as NBAD sclerotin (clear cuticle) when conjugated with β-alanin through the activity of ebony and black (responsible for metabolization of aspartate into β-alanin) enzymes. In bold are indicated the names of the key enzymes functionally characterized in Drosophila; just behind are indicated in italic type the name of the closest pea aphid homologue from the latest release of the pea aphid genome (v2.1). PO: enzymes from the phenoloxidase family.

In this paper, we aimed at pursuing our characterization of dopamine pathway in the context of reproductive mode switch in aphids. To achieve that, we first localized *pale* and *ddc* transcripts in embryos and then analyzed the expression of the key genes involved in the dopamine-derived pathways in embryos and heads of larvae under short and long days conditions. We finally targeted the *ddc* gene in fertilized eggs with CRISPR-Cas9 to generate mutant lineages and potentially analyze the precise role of dopamine in the transduction step of the photoperiodic response.

## Methods

### 1. Aphid rearing and production of sexuparae and virginoparae embryos

Stocks of virginoparae individuals of *Acyrthosiphon pisum* strain LSR1 (IAGC, 2010) are maintained on broad bean (*Vicia fabae*) plants in growth chambers at 18°C under long days conditions (16h of light, 8h of night), at low density (5 individuals per plant) to prevent the induction of winged morphs. In order to induce the production of sexual individuals, we used an already well-established protocol consisting in transferring individuals at precise developmental stages from long days to short days conditions (Le Trionnaire et al., 2009). Briefly, virginoparae L3 individuals were collected from the initial stock to produce in two different batches future sexuparae (under short days) or virginoparae (under long days). Future sexuparae are produced on broad bean at 18°C when placed at 12h of light (12h of night) while virginoparae are produced when maintained at 18°C under 16h of light (8h of night). Those individuals - when they become adult - represent generation 0 (G0). The embryos contained in those G0 are either future virginoparae (under long days) or sexuparae (under short days) and after having been laid on the plant until they reach adulthood, they correspond to generation 1 (G1). G1 virginoparae lay down virginoparae females at generation 2 (G2), while G1 sexuparae lay down sexual individuals (oviparous sexual females first and then males) at G2. This experimental design, as well as the developmental stages where biological material was collected for subsequent molecular analyzes (qPCR, RNA-seq and *in situ* hybridization) is summarized in Figure 2.

**Figure 2.**
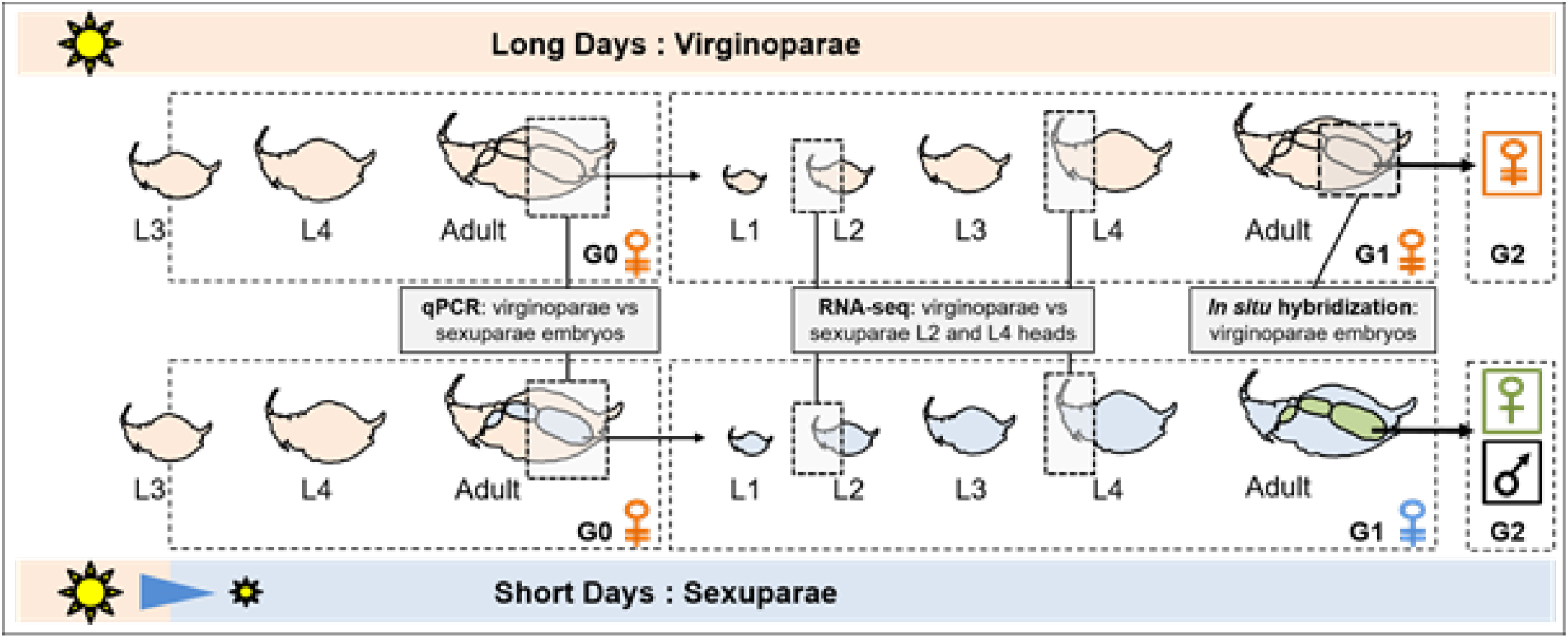
Experimental design used to collect biological material for molecular experiments. Briefly, under long photoperiod conditions only virginoparae females are produced (G0, G1 and G2). When transferred from long to short photoperiod at L3 stage, the embryos (G1) for these virginiparae females (G0) start to perceive changes in day length. Once born, these females continue to integrate this cumulative signal until it triggers the switch of the germline fate of their embryos (G2): these virginoparae females that produce sexual individuals (oviparous females and males) in their offspring are called “sexuparae”. In this study, we analysed the level of expression of dopamine pathway genes in virginoparae and sexuparae embryos (by qPCR) and L2 and L4 heads (extracted from RNA-seq data). *In situ* hybridization of *pale* and *ddc* transcripts was performed in virginoparae embryos chains.

### 2. Dopamine pathway gene annotation and PCR primers definition

In order to find pea aphid homologues for Drosophila genes functionally characterized for their involvement in dopamine pathway (as described in Figure 1), the amino acid sequences of *pale, ddc, vat1, prt, vmat, aaNAT, black, ebony, laccase* and *yellow* genes were retrieved from Flybase. A Blastp analysis was then performed on AphidBase (https://bipaa.genouest.org/is/aphidbase/) on the NCBI v2.1 annotation of *Acrythosiphon pisum* genome in order to find the closest homologues for these genes. These genes were then manually annotated to discriminate exon and intron sequences. Coding sequences for these 10 genes are available in Table S1 (https://doi.org/10.57745/XHUV1S). Primers for quantitative RT-PCR were then searched on these sequences using the Primer 3 software (http://primer3.ut.ee/) with default parameters and a maximum size of 150 nt for amplicon length. Primers sequences are listed in Table S2 (https://doi.org/10.57745/XHUV1S).

### 3. Quantitative RT-PCR analyzes

For Quantitative RT-PCR analysis of dopamine pathway gene expression levels, both virginoparae and sexuparae embryos (see above) were dissected from 25 adult G0 aphids (Figure 2). The 6/7 most developed embryos were dissected on ice for each individual, then pooled into liquid nitrogen and finally stored at - 80°C before RNA extraction. Approximately 180 embryos were collected in both conditions (short and long days). Three biological replicates were performed. Total RNAs were extracted using the RNeasy Plant Mini kit (Qiagen) according to manufacturer’s instructions. The optional DNase treatment was carried out with the RNase-Free DNase Set (Qiagen). RNA quality was checked and quantified by spectrophotometry (Nanodrop Technologies). Before reverse transcription, a second round of DNase digestion was added using the RQ1 RNase-free DNAse (Promega), in order to remove any putative residual DNA. One microgram of total RNA was used for cDNA synthesis using the Superscript III Reverse Transcriptase (Invitrogen) and a poly-T oligonucleotide primer (Promega) following the manufacturer’s instructions. The cDNAs were used for the quantitative PCR assay on a LightCycler 480 Real-Time PCR System using the SYBR Green I Master mix (Roche) according to the manufacturer’s instructions. A standard curve was performed for each gene (*pale, ddc, vat1, Vmat, prt, laccase, yellow, black, ebony, aaNAT*, and *RpL7* used as a reference gene) using serial dilutions of cDNA products in order to assess PCR primers efficiency. A dissociation curve was produced at the end of each run in order to check for non-specific amplifications. Each Q-RT-PCR was performed on three technical replicates. Thus, for each condition, data were obtained from three biological replicates with three technical replicates for a total of nine measurements per condition and gene. Relative quantification was performed using the standard curve method with normalization to the *A. pisum* ribosomal protein L7 transcript (*RpL7*, Nakabachi et al., 2005). In our conditions, *RpL7* expression was stable across samples (less than 1 Ct of difference between samples), thus confirming its invariant status. Absolute measures for each of the ten target genes (averaged among three replicates) were divided by the absolute measure of *RpL7* transcript. For each of the ten genes involved in the dopamine pathway, the normalized (to *RpL7* invariant gene) expression values (the average of the three technical replicates) calculated for embryos dissected from virginoparae and sexuparae mothers (three biological replicates) were compared using a one-way ANOVA. A p-value ≤ 0.05 was applied to identify differentially expressed genes between the two conditions. All raw data are accessible here: https://data.inrae.fr/privateurl.xhtml?token=d41cb434-97c3-400c-9d89-cb7c32299055 (Tagu et al. 2020).

### 4. RNA-seq data

In a previous study, we compared the transcriptomes of virginoparae (under long days or LD) and sexuparae (under short days or SD) head samples at two stages of larval development (L2-G1 and L4-G1, see Figure 2) using a custom-made cDNA microarray (Le Trionnaire et al., 2009). We recently used the exact same RNA samples (two biological replicates per time point) to perform RNA-seq analyses, for eight datasets (two for L2-G1 LD, two for L2-G1 SD, two for L4-G1 LD and two for L4-G1 SD). Raw reads from RNA-seq data were mapped onto the NCBI v2.1 annotation of Acrythosiphon pisum using STAR with default parameters (Dobin et al., 2015). Reads were then counted by genes using FeatureCounts (Liao et al., 2014) with default parameters. Counts normalisation and differential expression analyses have been performed using the scripts described in Law et al. (2016), which is based on EdgeR (Robinson et al., 2010). Genes with an adjusted p-value ≤ 0.05 were considered as differentially expressed between LD and SD conditions. From these data, the expression values and the p-value associated with the LD/SD comparisons were retrieved for the ten pea aphid homologues of Drosophila genes involved in the dopamine pathway (see above). Raw data are available on Sequence Read Archive (SRA) from NCBI under the SRP201439 accession number.

### 5. mRNA *in situ* hybridization

#### Riboprobe synthesis

Templates for riboprobes synthesis were amplified by RT-PCR, cloned and then transcribed into RNA. For this, total RNAs were extracted from adult parthenogenetic females (virginoparae) using the RNeasy Plant Mini kit (Qiagen). A DNase digestion step was carried out using RQ1 RNase-free DNAse (Promega) in order to remove any residual DNA. One microgram of total RNA was reversed transcribed with AMV Reverse Transcriptase (Promega) and a poly-T oligonucleotide primer (Promega) following the manufacturer’s instructions. The cDNA produced was used as a template for PCR amplification with specific primers for the two genes pale and ddc. Sequences of gene-specific primers and length of probes are given in Table S3 (https://doi.org/10.57745/XHUV1S). Amplified fragments were then cloned into StrataClone PCR Cloning Vector pSC-A-amp/kan (Stratagene) and sequenced in order to check for the identity and orientation of the inserted PCR fragment. Inserts containing the RNA polymerase promoters were obtained from the recombinant plasmids by PCR with universal primers. These PCR products (at least 500 ng per probe) were used as a template for synthesis of sense and antisense riboprobes using digoxigenin-labelled dNTPs (Dig RNA Labelling Mix (Roche)) and the appropriate RNA polymerase T7, T3 or SP6 (Roche). After synthesis, DNA was removed with RQ1 RNase-free DNAse treatment (Promega) and labelled riboprobes were purified with the RNeasy MinElute Cleanup kit (Qiagen). Riboprobe quality and quantity was checked on an agarose gel containing SybrSafe (Invitrogen) and quantified with Nanodrop (ThermoFicherScientific).

#### Whole-mount in situ hybridization on aphid ovaries

We adapted the *in situ* hybridization protocol previously described in Gallot et al. (2012). Ovaries containing the ovarioles of developing embryos were dissected from virginoparae adult individuals, maintained under long photoperiod (see Figure 2). For this, caudas were removed with clamps and ovaries chains were slightly disrupted from conjunctive tissues under a glass microscope slide within fixation solution (4% paraformaldehyde in PBS buffer). Dissected ovarioles were incubated with the probes of interest: 630 ng/ml for ddc and 1000 ng/ml for pale in the same conditions as described in Gallot et al. (2012). Detection was performed with anti-DIG-alkaline phosphatase (AP) Fab fragments (Roche) diluted 1:2000 in blocking solution. Signal was revealed with 4 μL of NitroBlue Tetrazolium/5-Bromo-4-Chloro-3-Indolyl Phosphate (NBT/BCIP) Stock Solution (Roche)/ml of AP reaction buffer. Ovarioles were then observed under a microscope Nikon 90i connected to a Nikon type DS-Ri1 camera to allow images capture. Controls were performed using the corresponding sense probes.

### 6. CRIPSR-Cas9 editing of *ddc* gene

In order to analyze the function of ddc gene in the context of aphid reproductive mode switch, we aimed at generating stable mutant lineages for this gene. The step-by-step protocol for CRISPR-Cas9 mutagenesis in the pea aphid is detailed in Le Trionnaire et al. (2019). Briefly, we designed two single guide RNAs (sg1 and sg2) predicted to target the fifth exon of *ddc* gene using the CRISPOR software (Figure 7a). We then performed *in vitro* cleavage assays to confirm the efficiency of these guide RNAs to cut - when complexed to Cas9 - a PCR product from *ddc* genomic sequence boarding the fifth exon (Figure 7b). The primers used to amplify these genomic regions are the following: F-ACTTTGGTGGCGTTGTTG and R-ggacggaggcacagactaag. We then induced the production of sexual females from L9Ms10 clone and males from L7Tp23 clone to obtain fertilized eggs and inject them with a sg1/sg2/Cas9 mix and waited until melanisation before placing them at 4°C for 85 days for obligate diapause.

## Results

### Dopamine pathway genes are expressed in the heads of sexuparae and virginoparae

In the past, our group demonstrated a decrease of the steady-state level of the *pale* and *ddc* mRNAs in the heads of pea aphid sexuparae under short days (SD) when compared with virginoparae aphids reared under long days (LD). This down regulation was detected at the third larval stage (L3) of sexuparae (Gallot et al., 2010). These two genes encode two key enzymes involved in the formation of dopamine from L-tyrosine. In order to test whether the various downstream genes of dopamine pathway signaling (Figure 1) were regulated, we used transcriptomic data generated in aphid heads from LD (virginoparae) and SD (sexuparae) reared aphids at L2 and L4 larval stages. We performed RNA-seq on samples already used for cDNA arrays (Le Trionnaire et al. 2009) and extracted the expression information specifically for the dopamine pathway genes (Figure 3). Statistical analyses revealed that *ddc* and *pale* transcript were significantly down regulated under SD conditions at L2 and L4 stages thus confirming the trend observed at L3 stage. Regarding the genes involved in the synaptic release of dopamine, only *prt* transcript was down regulated at L4 stage under SD conditions while *vat1* and *vmat* were not regulated by photoperiod shortening. All the genes involved in the use of dopamine-derived molecules as an uptake for cuticle melanization and sclerotization were regulated between LD and SD conditions. More precisely, *aaNAT, black, ebony* and *yellow* were significantly down-regulated under SD conditions at L2 and L4 stages while *laccase* transcript was up-regulated at L2 stage and down-regulated at L4 stage. These data thus indicate that in sexuparae heads, genes involved in cuticle sclerotisation and melanisation are down-regulated under SD conditions while it is not the case for genes involved in synaptic dopamine transport.

**Figure 3.**
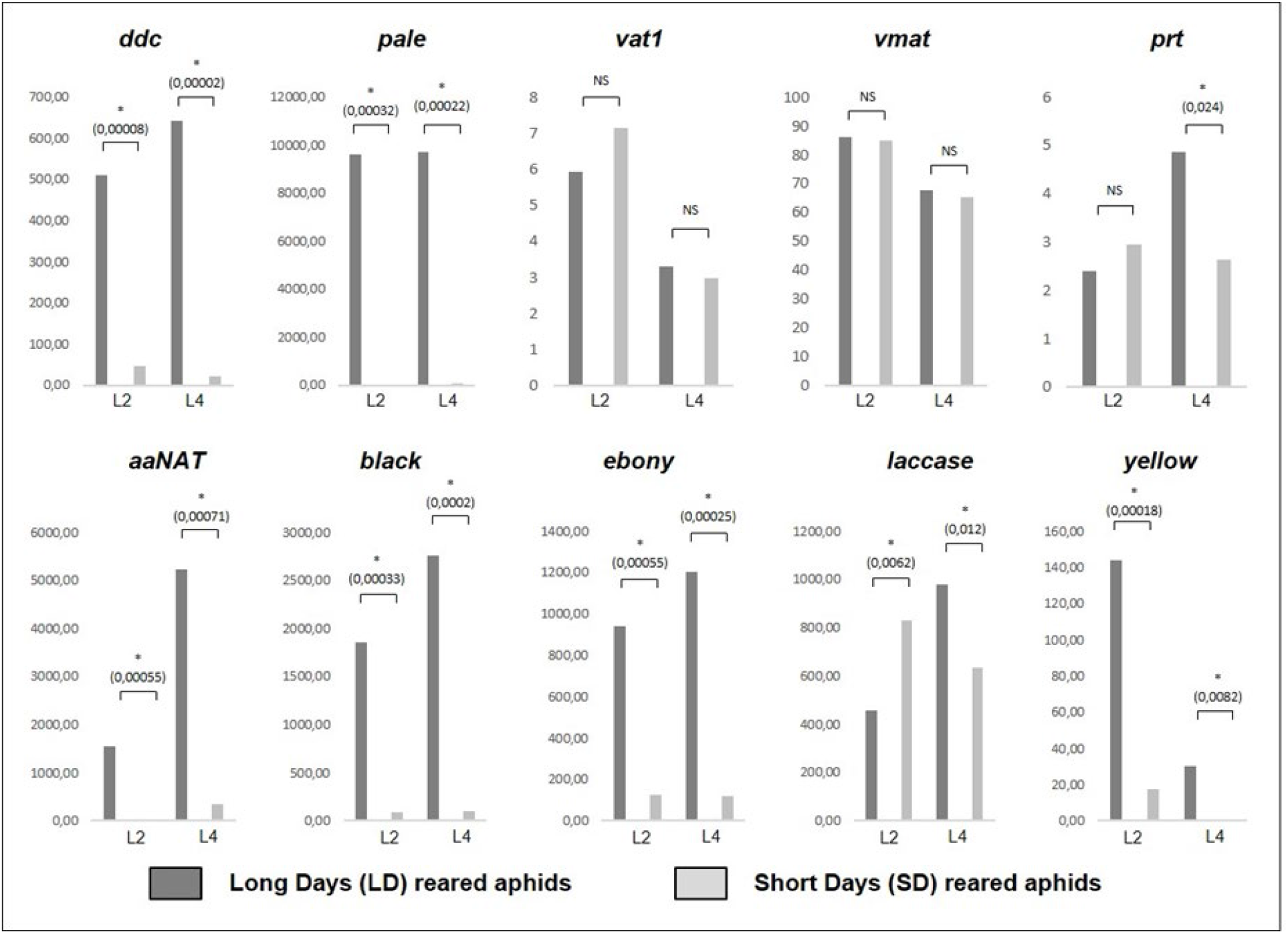
Expression levels of dopamine pathway genes in L2 and L4 larvae heads of long day (LD) and short day-reared aphids. Based on a previous microarray analysis of the transcriptomic response in the heads of aphids when submitted to photoperiod shortening (Le Trionnaire et al., 2009), the same RNA samples were used for RNA-sequencing. The level of expression (expressed as RPKM) of dopamine pathway genes was extracted from this dataset and statistically analyzed using EdegR package to identify differentially expressed genes between the two photoperiod conditions (the p-value of the test is indicated between brackets). *: Significant, NS: Non-Significant.

### Localization of *pale* and *ddc* transcripts in virginoparae embryos

In order to test whether *pale* and *ddc* are also expressed prenatally, we performed in situ localization of the pea aphid mRNAs in the embryos (and ovarioles) of adult aphids, just before larvae birth. *pale* mRNAs were detected in one early stage of embryogenesis (stage 1, as defined by Miura et al., 2003), in a very restricted zone (Figure 4). This pattern was reproducible, but difficult to interpret. As it was not detectable in subsequent early stages, we propose the hypothesis that it might correspond to a residual signal of maternal pale mRNAs. *pale* mRNAs were also detected at later embryogenesis stages (from stage 13 to 17/18), at the anterior part of the embryo, probably in the protocerebrum, and more particularly in discrete pairs of cells that might correspond to neurosecretory cells. We could however not clearly assign the specific type of neurosecretory cells they could correspond to (Steel, 1977). No labeling was detected in the downstream ganglion chain, but a lateral labeling was detectable within unknown structures that might be mushroom bodies (Kollmann et al., 2011). *ddc* mRNAs were detected in the latest stages (from stages 16 to 18) of embryogenesis (Figure 5). Labeled cells were located within the anterior part of the embryo, in neuronal cells of the protocerebrum that are likely to correspond to neurosecretory Cells from Group I, II, III or IV, which are located in a median position of the brain (Steel, 1977). Labeled cells were also detectable posteriorly, along the ganglion chain of the nervous system. Thus, *pale* and *ddc* genes are both expressed in the embryonic central nervous system but have distinct spatial patterns of expression. As expected, no labeled structures were detected in control samples hybridized with sense probes (Figures 4 and 5).

**Figure 4.**
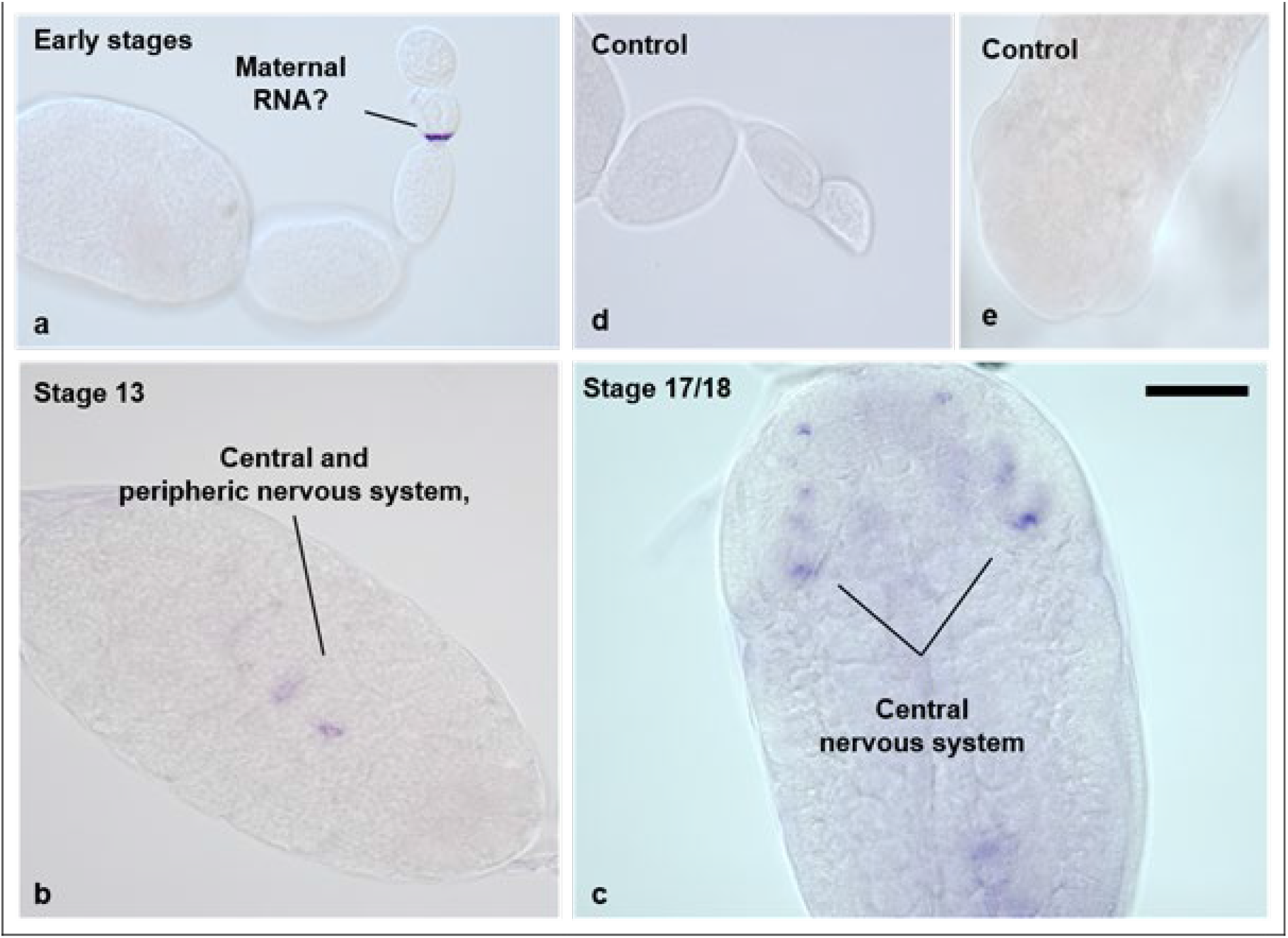
In situ hybridization of pale transcripts within virginoparae embryos. Ovaries were dissected from virginoparae morphs and hybridized with antisense (a, b and c) or sense (d and e) probe.

**Figure 5.**
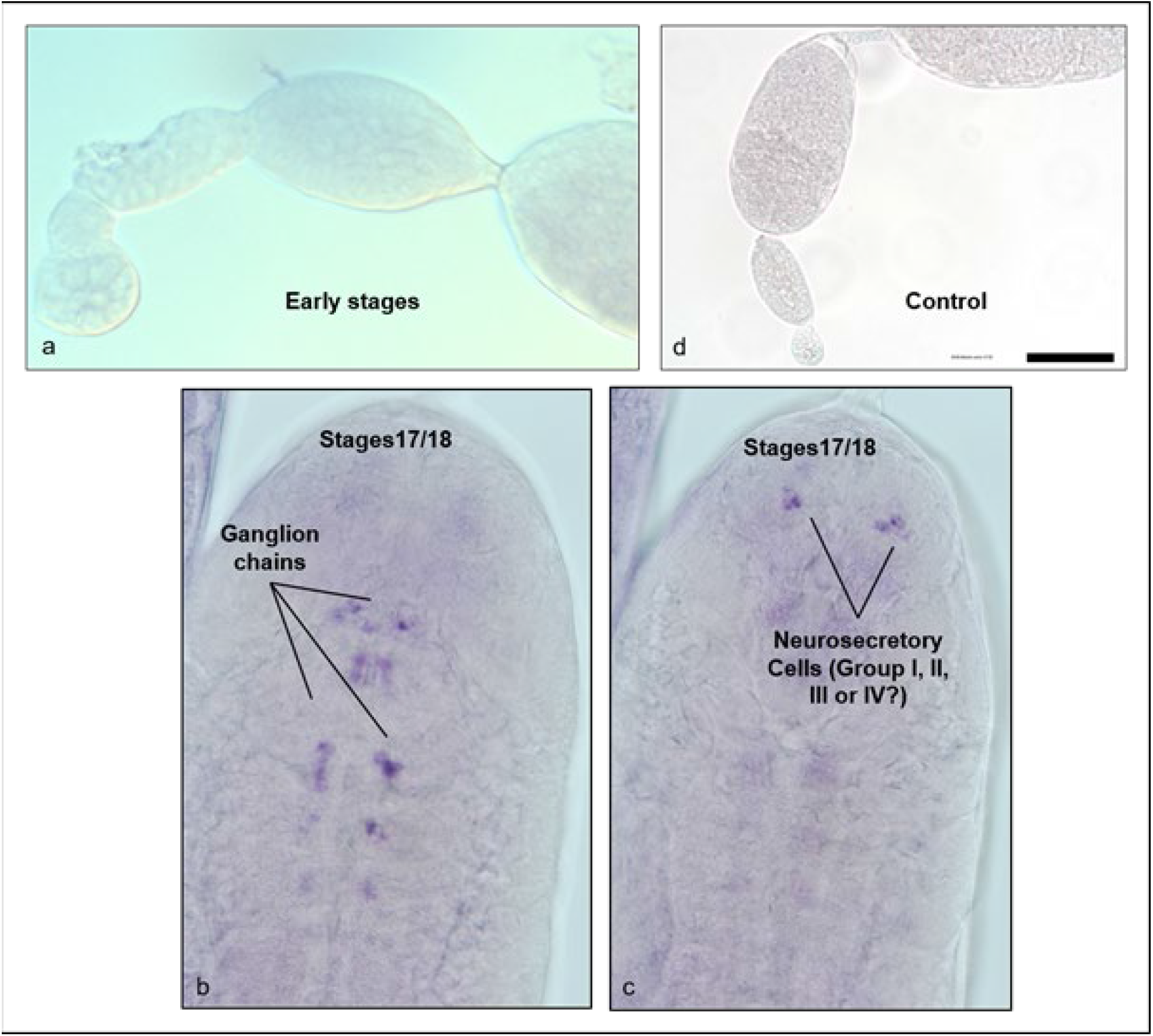
In situ hybridization of ddc transcripts within virginoparae embryos. Ovaries were dissected from virginoparae morphs and hybridized with antisense (a, b and c) or sense (d) probe.

### Dopamine pathway genes expression in sexuparae and virginoparae embryos

We checked whether the dopamine pathway genes were differentially expressed in embryos between LD and SD photoperiodic regime (Figure 2). We compared by qRT-PCR the expression of the corresponding mRNAs between late stages (see methods) of sexuparae and virginoparae embryos (Figure 6). *vat1, vmat* and *prt* which are genes involved in the synaptic release of dopamine were consistently not regulated (with a p-value close to 1) between the two types of embryos. Then *ddc* and *pale* appeared to be more expressed in virginoparae embryos compared with sexuaparae embryos, but because of an important variability between replicates, these differences were not significant. Regarding the genes involved in cuticle melanization and sclerotization, laccase and yellow were significantly down regulated in sexuparae embryos. *aaNAT, black* and *ebony* also appeared to be more expressed in virginoparae embryos. The observed differences, while not in all cases statistically significant (p < 0.05), were in a consistent direction (down regulation in sexuparae). These analyses show that cuticle-related genes (especially laccase and yellow) are already down-regulated in SD-reared embryos while the expression of the others are probably in the process of being altered to reach the patterns observed after birth at L2 and L4 stages.

**Figure 6.**
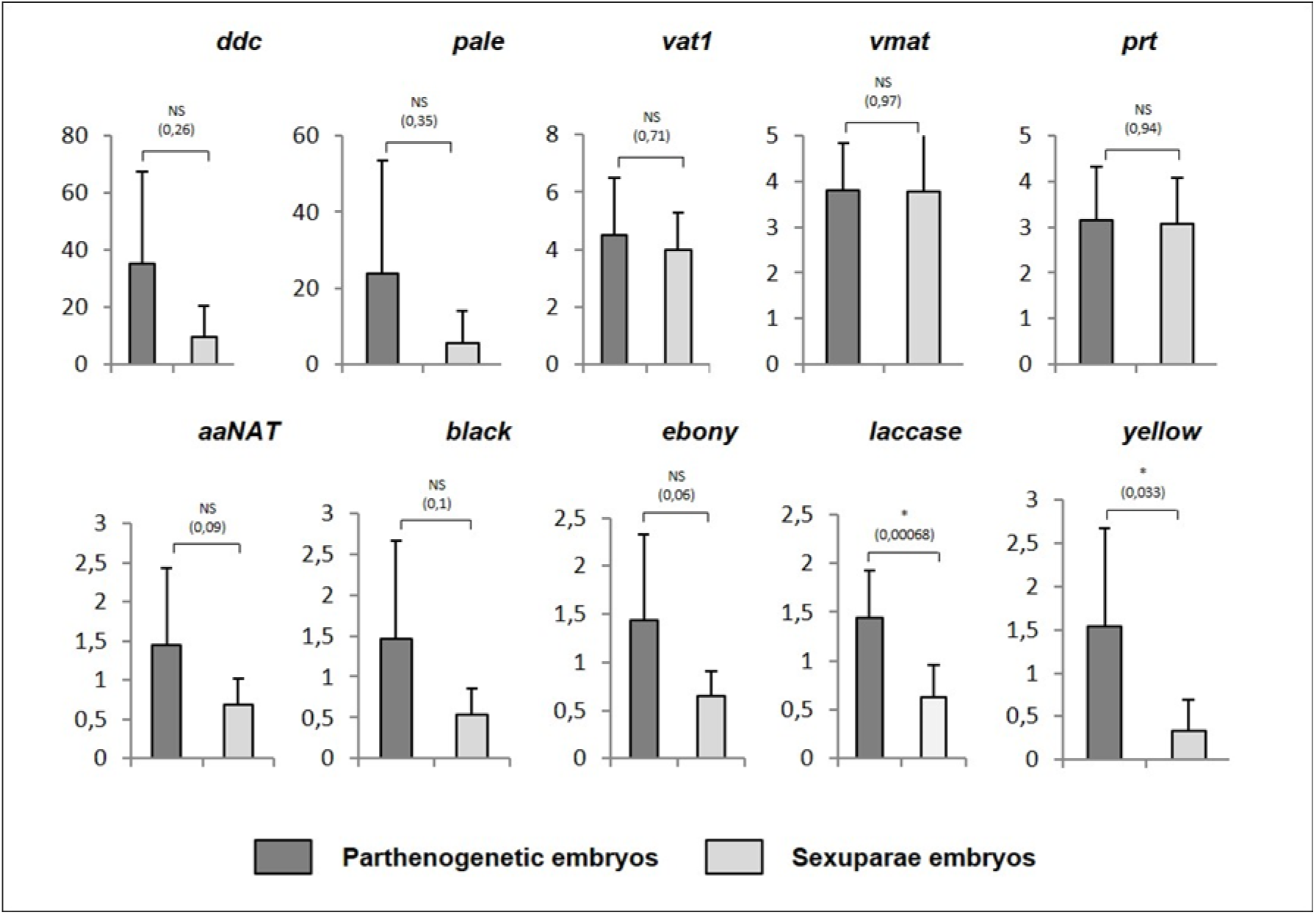
Expression levels of dopamine pathway genes in virginoparae and sexuparae embryos. Most developed embryos from adult aphids placed under long days (virginoparae embryos) and short days (sexuparae embryos) were collected for the quantification of the level of expression of dopamine pathway genes. Relative quantification was performed using RpL7 as an invariant gene. A t-test was performed to estimate if the genes were significantly regulated between the two conditions (the p-value of the test is indicated between brackets). *: Significant, NS: Non-Significant.

### CRISPR-Cas9 mutagenesis of ddc gene in fertilized eggs

Based on a recent protocol of CRISPR-Cas9 mutagenesis developed in the pea aphid (Le Trionnaire et al., 2019), we aimed at generating mutant lineages for *ddc* gene. We injected fertilized eggs with two single guide RNAs designed to target the fourth exon and some recombinant Cas9 protein (Figure 7a). These guides were validated in vitro prior to injection in order to maximise the possibility to generate genome editing events (Figure 7b). Finally, we injected 851 fertilized eggs less than 4 hours after being laid on the plant by the sexual females. Among them, 470 (55%) of the eggs were damaged by the micro-injection procedure, a percentage closed to what we observed in previous experiments. Among the remaining eggs, 84 (10%) completed melanisation, indicating their survival. Surprisingly, 297 eggs (35%) appeared to be intact (not damaged by the injection) but did not complete melanisation (Figure 7c). Indeed, melanisation in aphid eggs is associated with a gradual transition from green to black colour and usually lasts for 5 days, which corresponds to the early steps of embryogenesis until melanin synthesis is completed. The colour defaults we observed ranged from green-spotted dark eggs, black-spotted green eggs and finally eggs that remained almost entirely green. As these colour patterns are absent in non-injected or water-injected eggs, these observations are likely to correspond to a melanisation default phenotype due to the genome editing of *ddc* gene. In order to confirm that CRISPR-Cas9 system had effectively generated mutations, we collected 63 eggs showing this various colour phenotypes (Figure 7d). We then extracted the DNA of each of these eggs and amplified by PCR the *ddc* genomic region. A gel electrophoresis analysis showed that at least 63% (40/63) of the eggs displayed the wild-type band but also some additional bands of smaller size. These products are the result of the combined action of the two sgRNAs that provoked some large deletion events around the two target sites. This analysis thus revealed that these eggs with melanisation defaults are mutated for the *ddc* gene. A focus (Figure 7e) on nine eggs with colour defaults clearly shows the presence of extra bands and their absence in non-injected eggs (NI). This genome editing experiment suggests that *ddc* gene in the pea aphid is involved in cuticle melanisation. Nevertheless, this phenotype is lethal since eggs with incomplete melanisation did not survive. Consequently, those eggs could not hatch and we were unable to generate stable mutant lineages for *ddc* gene and consequently we could not to test the effect of a *ddc* knock out on the reproductive mode switch of aphids.

**Figure 7.**
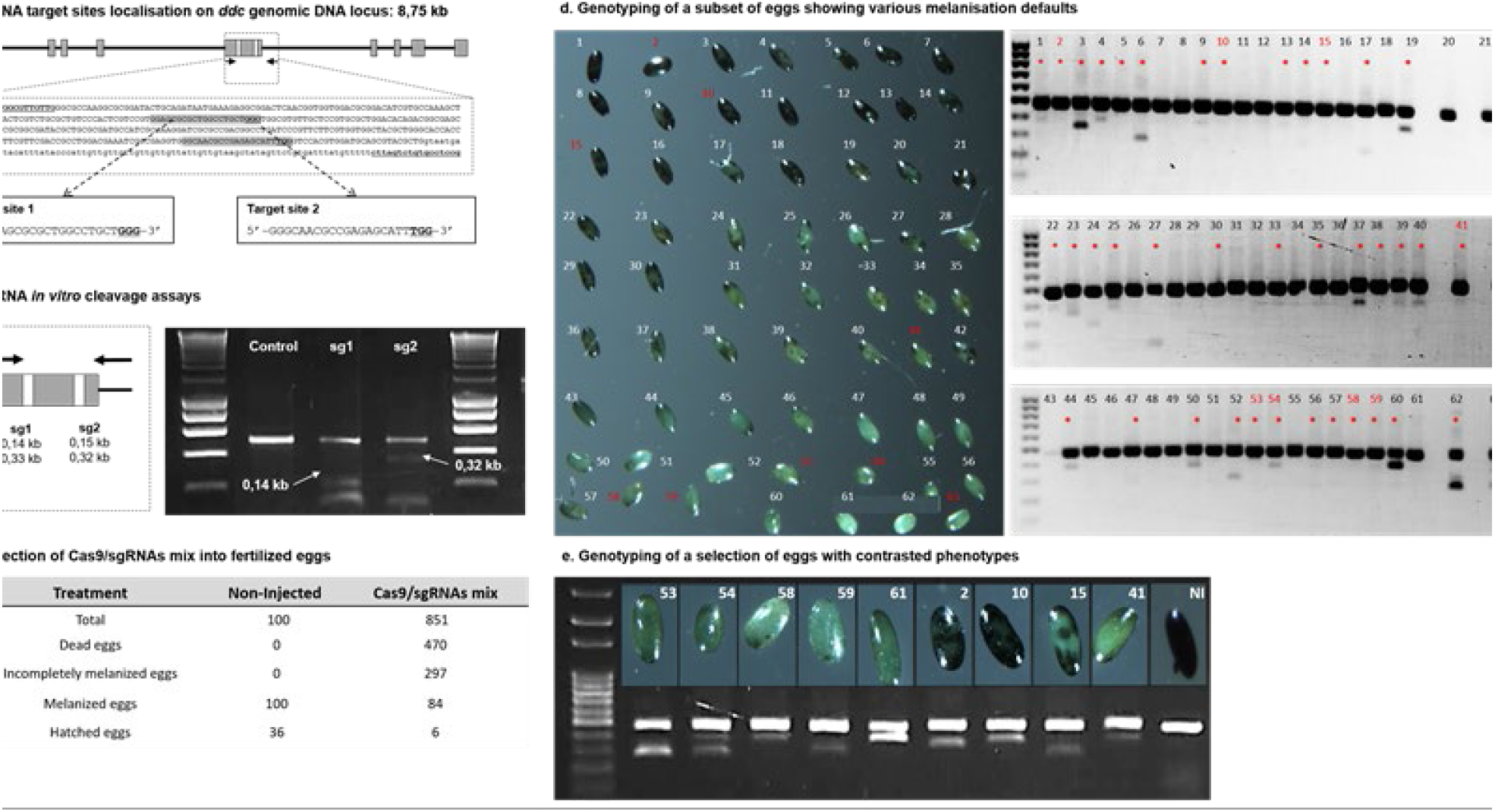
CRISPR-Cas9 editing of ddc gene and phenotypic defaults observed in injected fertilized eggs. a) The ddc coding sequence comprises 9 exons that are spread over a 8,75 kb genomic DNA region. Single guide RNAs were defined to target two regions (target site 1 and 2) within the 5th exon of the gene. On the 5th exon sequence are underlined the Forward and Reverse primers used to amplify the corresponding genomic DNA. b) In vitro cleavage assay were performed by mixing each sgRNA, the cas9 protein and the PCR product of the 5th exon. The sizes of the expected cleavage products are indicated with white arrows. c) This table compiles the results from the micro-injection within fertilized eggs of a sg1/sg2/Cas9 mix in terms of dead eggs, melanised eggs but also eggs with incomplete melanisation. Post-diapause hatching rates are also indicated, including the non-injected control eggs. d) 63 eggs with melanisation defaults were collected and their ddc genomic location was amplified. Gel electrophoresis analysis reveals the presence of extra bands that are the product of the combined action of the two sgRNAs and confirms the efficiency of genome editing (red stars). e) A focus on nine mutated eggs and one non-injected egg confirms the efficiency of CRISR-Cas9 system to generate contrasted melanisation phenotypes.

## Discussion

Dopamine is a central regulator of various pathways in insects, including synaptic function and cuticle structure. Previous works proposed the hypothesis that dopamine could be involved in the regulation of the plasticity of the reproductive mode in aphids (Le Trionnaire et al., 2009 and Gallot et al., 2010). This was based on the differential expression between long days (LD) and short days (SD) conditions of genes involved in dopamine synthesis as well as dopamine conjugation with other components to form cuticle polymers. Dopamine pathway (or more generally catecholamine pathways) has been demonstrated to regulate the phase polyphenism in locust, another case of discrete phenotypic plasticity where solitary and gregarious morphs alternate upon population density changes (reviewed in Wang and Kang, 2014). More precisely, it was shown that three genes involved in dopamine synthesis and synaptic release (*pale, henna* and *vat1*) were differentially expressed between the alternative morphs of Locusta migratoria (Ma et al., 2011). Artificial modification of dopamine levels clearly demonstrated the role of this pathway in the control of phase polyphenism. The switch of reproductive mode in aphids being another striking case of insect polyphenism, we made the hypothesis that dopamine might be a good candidate as a potential signaling pathway linking photoperiod shortening perception and asexual to sexual embryogenesis transition.

In this work, we analysed the expression levels within A. pisum embryos and heads of larvae of most of the genes involved in dopamine-related pathways. When comparing RNA-seq expression levels of these genes in head tissues, it clearly appeared that the genes coding for key enzymes involved in dopamine synthesis (*pale* and *ddc*) and the genes involved in cuticle structure changes (*aaNAT, black, ebony, laccase* and *yellow*) were down-regulated in SD-reared aphids (sexuparae individuals). On the contrary, genes involved in dopamine transport and release in synapses (*vat1, vmat* and *prt*) were globally not regulated, apart from the *prt* homologue that was down-regulated under SD conditions but only at a specific stage (L4). Altogether, these data confirm previous observations that photoperiod shortening in sexuparae individuals may result in cuticle structure modifications triggered by a potential decrease in dopamine synthesis levels (Le Trionnaire et al., 2009). We then tested whether these changes were initiated prenatally as a result of an early perception of photoperiod shortening in embryos. We compared the expression levels of these genes in LD-reared embryos (future vriginoparae) and SD-reared embryos (future sexuparae). Some of these genes – including *pale, ddc, aaNAT, black* and *ebony* - showed a tendency of being less expressed (although not significant) in sexuparae embryos while some others were significantly down-regulated (e.g. *laccase* and *yellow*) in comparison with virginoparae embryos. Again, no differences in expression could be observed for genes involved in dopamine release in synapse. Altogether, these data suggest that the photoperiod decrease might affect the level of expression of dopamine synthesis genes and somehow reduce dopamine levels in embryos and latter in heads of larvae which in turn would promote changes in cuticle structure, as confirmed by similar changes in the expression of genes coding for enzymes involved in this process. Regarding the absence of regulation of genes coding for dopamine release in synapses, this does not rule out the possibility that dopamine might serve as an intermediate signaling neurotransmitter linking photoperiod and embryos fate change, assuming that these type of transporters could possibly be ubiquitously expressed whether in LD and SD-reared embryos or in LD and SD-reared heads. Although our data cannot help concluding on that point, we can still speculate that dopamine levels might start to decrease prenatally following an early perception of SD photoperiodic regime and then continue to do so after birth until the required number of short days (Lees, 1989) necessary to induce the switch in embryos is reached. A time-course experiment of dopamine concentration measurement in LD and SD reared embryos and heads, as well as later in asexual and sexual embryos might certainly help deciphering the role of dopamine as a signaling molecule during the whole process of reproductive mode switch.

Transcripts coding for key dopamine synthesis enzymes (*pale* and *ddc*) are expressed and specifically localized in embryos, more precisely in paired structures and cells in the central nervous system and associated neuronal ganglions. However, due to the lack of a precise description of the anatomy of aphid embryonic central nervous system, it is difficult to identify with accuracy the cell types expressing those transcripts. We detected a *ddc* mRNA signal in cells that could correspond to neurosecretory cells described in adult aphids (Steel, 1977). These labelled cells putatively belonging to Group I, II, III or IV of neurosecretory cells. Some of them consist in five neurons located in the pars intercerebralis of the protocerebrum (Group I), and one to two cells at the posterior part of the protocerebrum (Group IV). Interestingly, Group I cells have been showed to be involved in the photoperiodic response in the aphid Megoura viciae (Steel and Lees, 1977). Specific lesions of these cells abolished the response to the photoperiod. However, the nature of the material secreted by these cells is not known. They resemble neurosecretory cells of other insects such as D. melanogaster known to produce insulin peptides (Cao and Brown, 2001). There is nevertheless no clear evidence that *ddc* and *pale* positive cells/neurons are these neurosecretory cells and a more detailed analysis of the localization of *ddc* mRNAs in the central nervous system of aphid embryos and brain structures are required, including immuno-localisation of the corresponding protein. Finally, these data allow us to conclude that embryos already express dopamine synthesis genes in dedicated nervous structures from the brain and the central nervous system and that photoperiod somehow affect their level of expression.

The down-regulation of dopamine synthesis pathway observed in the embryos is correlated with cuticle structure modifications. The hypothesis we made is that the head cuticle of sexuparae is less sclerotized (Gallot et al., 2010, this study) and less melanized (this study) than in virginoparae. Whether this fact is a cause or a consequence of the production of alternative reproductive morphs in aphids in not solved yet. In one hand, cuticle structure could modify brain light perception by the sexuparae, but whether or not this has a direct incidence on the plasticity is not known. On the other hand, sexuparae have a very similar morphology to the virginoparae, but to our knowledge, no deep scanning of possible differences has been performed (except for the type of embryos). Cuticle structure could be one of these general phenotypical differences and probably needs to be investigated into more details.

In order to test the possibility that dopamine might be a signaling intermediate between photoperiod shortening and embryos fate change, we aimed at knocking-out *ddc* gene to disturb dopamine synthesis. In this context, an expectable phenotype was that knocked-out lineages might produce sexual individuals directly after hatching even under LD conditions. We did not succeed to knock-down the *ddc* transcript in the pea aphid by RNAi (data not shown), probably because of the specific expression of these genes in the brain and central nervous system which are tissues/organs difficult to target by RNAi in aphids. We thus moved to a knock-out approach to edit the *ddc* gene with CRISPR-Cas9 by injecting fertilized eggs with some Cas9 protein together with a pair of single guide RNAs designed to target one specific exon of the ddc gene. Surprisingly, the mortality of injected eggs was particularly high in comparison with previous experiments of gene editing on another candidate (Le Trionnaire et al., 2019). Indeed, only a few eggs reached complete melanization while the remaining proportion of the eggs (that were not damaged by the injection procedure) displayed various patterns of melanization, from nearly absent to almost complete. It is known in Drosophila that *ddc* (alongside with *pale*) is required for melanin synthesis (True et al., 1999). Molecular analyzes revealed the presence of mutations in those eggs so that mutants for *ddc* gene display some body color defects, especially a clearer color due to the absence of melanin. The phenotype observed in aphid eggs thus resembles the Drosophila phenotype. This indicates that CRISPR-Cas9 system generated knock-out of the *ddc* gene in eggs that show an altered melanin synthesis in various locations. It is likely that these eggs harbor distinct somatic mutations in various embryonic tissues responsible for a mosaicism materialized by the variety of color patterns observed. Since the addition of melanin at the surface of the egg is necessary to protect the embryo against winter environmental conditions this explains why these eggs could not hatch after diapause and why this mutation had a lethal effect in our conditions. This experiment thus revealed that *ddc* gene was involved in melanization in aphids. Nevertheless, the lethal effect of the *ddc* knockout on egg viability did not allow us to investigate the role of this gene in the response to photoperiod since we could not establish stable mutant lineages. Nevertheless, this study represents so far the first mutant phenotype ever generated on the aphid model with CRISPR-Cas9. Despite not providing a straightforward answer on the role of *ddc* during reproductive mode switch in aphids, these results confirm the efficiency and reproducibility of the targeted mutagenesis protocol we developed in the pea aphid (Le Trionnaire et al., 2019) and that will be useful for the aphid community.

In conclusion, further functional analyses are required to dissect precisely the specific role of dopamine pathway in the regulation of reproductive polyphenism in aphids. Nevertheless, our study already demonstrated that various dopamine pathways genes were regulated prenatally between LD and SD reared embryos. This trend was confirmed in larvae where several of these genes were differentially expressed in the head between asexual and sexuparae (sexual-producers) individuals. This suggests that the setting of the genetic programs involved in the production of seasonal alternative reproductive morphs in the pea aphid takes place early in this trans-generational process, which is correlated with the development and differentiation of neuronal structures and cells within the embryos. Our results also indicate that photoperiod shortening is correlated with a reduction in the levels of expression of enzymes involved in dopamine synthesis that might affect cuticle structure. Our data nevertheless did not help us to suggest that dopamine might also act as signaling molecule linking photoperiod and embryos fate switch. The melanization defaults observed for *ddc*-mutated eggs by CRISPR-Cas9 eventually confirmed that dopamine has a major role in aphid cuticle composition and synthesis.

## Acknowledgements

We appreciated discussion with Dr. Sylvia Anton and Dr. Christophe Gadenne (INRAE, UMR Igepp, Rennes, France). Version 4 of this preprint has been peer-reviewed and recommended by Peer Community In Zoology (https://doi.org/10.24072/pci.zool.100013).

## Data, scripts and codes availability

RNA-Seq data: Sequence Read Archive (SRA) from NCBI, SRP201439 accession number.Q-RT-PCR data: https://data.inrae.fr/privateurl.xhtml?token=d41cb434-97c3-400c-9d89-cb7c32299055 (Tagu et al. 2020).

## Supplementary material

Supplementary material are available online: https://data.inrae.fr/dataset.xhtml?persistentId=doi:10.57745/XHUV1S.

## References

Andersen SO (2010) Insect cuticular sclerotization: a review. Insect Biochemistry and Molecular Biology, 40, 166–178. http://dx.doi.org/10.1016/j.ibmb.2009.10.007

Arakane Y, Dittmer NT, Tomoyasu Y, Kramer KJ, Muthukrishnan S, Beeman RW, Kanost MR (2010) Identification, mRNA expression and functional analysis of several yellow family genes in Tribolium castaneum. Insect Biochemistry and Molecular Biology, 40, 259–266. https://doi.org/10.1016/j.ibmb.2010.01.012

Arakane Y, Lomakin J, Beeman RW, Muthukrishnan S, Gehrke SH, Kanost MR, Kramer KJ (2009) Molecular and functional analyses of amino acid decarboxylases involved in cuticle tanning in Tribolium castaneum. Journal of Biological Chemistry, 284, 16584–16594. https://doi.org/10.1074/jbc.M901629200

Brey P, Ohayon H, Lesourd M, Castex H, Roucache J, Latge J (1985) Ultrastructure and chemical composition of the outer layers of the cuticle of the pea aphid Acyrthosiphon pisum (Harris). Comparative Biochemistry and Physiology Part A: Physiology, 82, 401–411. http://erepository.uonbi.ac.ke:8080/xmlui/handle/123456789/33962

Cao C, Brown MR (2001) Localization of an insulin-like peptide in brains of two flies. Cell and Tissue Research, 304, 317–321. https://doi.org/10.1007/s004410100367

Consortium IAG (2010) Genome sequence of the pea aphid Acyrthosiphon pisum. PLoS Biology, 8, e1000313. https://doi.org/10.1371/journal.pbio.1000313

Cortes T, Tagu D, Simon JC, Moya A, Martinez-Torres D (2008) Sex versus parthenogenesis: a transcriptomic approach of photoperiod response in the model aphid Acyrthosiphon pisum (Hemiptera: Aphididae). Gene, 408, 146–156. https://doi.org/10.1016/j.gene.2007.10.030

Dobin A, Gingeras TR (2015) Mapping RNA-seq reads with STAR. Current Protocols in Bioinformatics, 51. https://doi.org/10.1002/0471250953.bi1114s51

Febvay G, Pageaux JF, Bonnot G (1992) Lipid composition of the pea aphid, Acyrthosiphon pisum (Harris)(Homoptera: Aphididae), reared on host plant and on artificial media. Archives of Insect Biochemistry and Physiology, 21, 103–118. https://doi.org/10.1002/arch.940210204

Gallot A, Rispe C, Leterme N, Gauthier JP, Jaubert-Possamai S, Tagu, D (2010) Cuticular proteins and seasonal photoperiodism in aphids. Insect Biochemistry and Molecular Biology, 40, 235–240. https://doi.org/10.1016/j.ibmb.2009.12.001

Gallot A, Shigenobu S, Hashiyama T, Jaubert-Possamai S, Tagu D (2012) Sexual and asexual oogenesis require the expression of unique and shared sets of genes in the insect Acyrthosiphon pisum. BMC Genomics, 13, 76. https://doi.org/10.1186/1471-2164-13-76

Jaquiéry J, Peccoud J, Ouisse T, Legeai F, Prunier-Leterme N, Gouin A, Nouhaud P, Brisson JA, Bickel R, Purandare S (2018) Disentangling the causes for faster-X evolution in aphids. Genome Biology and Evolution, 10, 507–520. https://doi.org/10.1093/gbe/evy015

Kollmann M, Minoli S, Bonhomme J, Homberg U, Schachtner J, Tagu D, Anton S (2011) Revisiting the anatomy of the central nervous system of a hemimetabolous model insect species: the pea aphid Acyrthosiphon pisum. Cell and Tissue Research, 343, 343–355. https://doi.org/10.1007/s00441-010-1099-9

Law CW, Alhamdoosh M, Su S, Dong X, Tian L, Smyth GK, Ritchie ME (2016) RNA-seq analysis is easy as 1-2-3 with limma, Glimma and edgeR. F1000Research, 5. https://doi.org/10.12688/f1000research.9005.3

Lees AD (1964) The location of the photoperiodic receptors in the aphid Megoura viciae Buckton. Journal of Experimental Biology, 41, 119–133. https://doi.org/10.1242/jeb.41.1.119

Lees AD (1989) The photoperiodic responses and phenology of an English strain of the pea aphid Acyrthosiphon pisum. Ecological Entomology, 14, 69–78. https://doi.org/10.1111/j.1365-2311.1989.tb00755.x

Legeai F, Shigenobu S, Gauthier JP, Colbourne J, Rispe C, Collin O, Richards S, Wilson AC, Murphy T, Tagu, D (2010) AphidBase: a centralized bioinformatic resource for annotation of the pea aphid genome. Insect Molecular Biology, 19, 5–12. https://doi.org/10.1111/j.1365-2583.2009.00930.x

Le Trionnaire G, Francis F, Jaubert-Possamai S, Bonhomme J, De Pauw E, Gauthier JP, Haubruge E, Legeai F, Prunier-Leterme N, Simon JC (2009) Transcriptomic and proteomic analyses of seasonal photoperiodism in the pea aphid. BMC Genomics, 10, 456. https://doi.org/10.1186/1471-2164-10-456

Le Trionnaire G, Tanguy S, Hudaverdian S, Gleonnec F, Richard G, Cayrol B, Monsion B, Pichon E, Deshoux M, Webster C (2019) An integrated protocol for targeted mutagenesis with CRISPR-Cas9 system in the pea aphid. Insect Biochemistry and Molecular Biology, 110, 34–44. https://doi.org/10.1016/j.ibmb.2019.04.016

Le Trionnaire G, Wucher V, Tagu D (2013) Genome expression control during the photoperiodic response of aphids. Physiological Entomology, 38, 117–125. https://doi.org/10.1111/phen.12021

Liao Y, Smyth GK, Shi W (2014) featureCounts: an efficient general purpose program for assigning sequence reads to genomic features. Bioinformatics, 30, 923–930. https://doi.org/10.1093/bioinformatics/btt656

Ma Z, Guo W, Guo X, Wang X, Kang L (2011) Modulation of behavioral phase changes of the migratory locust by the catecholamine metabolic pathway. Proceedings of the National Academy of Sciences USA, 108, 3882–3887. https://doi.org/10.1073/pnas.1015098108

Miura T, Braendle C, Shingleton A, Sisk G, Kambhampati S, Stern DL (2003) A comparison of parthenogenetic and sexual embryogenesis of the pea aphid Acyrthosiphon pisum (Hemiptera: Aphidoidea). Journal of Experimental Zoology Part B: Molecular and Developmental Evolution, 295, 59–81. https://doi.org/10.1002/jez.b.3

Nakabachi A, Shigenobu S, Sakazume N, Shiraki T, Hayashizaki Y, Carninci P, Ishikawa H, Kudo T, Fukatsu T (2005) Transcriptome analysis of the aphid bacteriocyte, the symbiotic host cell that harbors an endocellular mutualistic bacterium, Buchnera. Proceedings of the National Academy of Sciences USA, 102, 5477–5482. https://doi.org/10.1073/pnas.0409034102

Ogawa K, Miura T (2014) Aphid polyphenisms: trans-generational developmental regulation through viviparity. Frontiers in Physiology, 5, 1. https://doi.org/10.3389/fphys.2014.00001

Robinson MD, McCarthy DJ, Smyth GK (2010) edgeR: a Bioconductor package for differential expression analysis of digital gene expression data. Bioinformatics, 26, 139–140. https://doi.org/10.1093/bioinformatics/btp616

Steel CGH (1977) The neurosecretory system in the aphid Megoura viciae, with reference to unusual features associated with long distance transport of neurosecretion. General and Comparative Endocrinology, 31, 307–322. https://doi.org/10.1016/0016-6480(77)90095-8

Steel C, Lees A (1977) The role of neurosecretion in the photoperiodic control of polymorphism in the aphid Megoura viciae. Journal of Experimental Biology, 67, 117–135. https://doi.org/10.1242/jeb.67.1.117

Tagu D, Le Trionnaire G, Tanguy S, Gauthier JP, Huynh JR (2014) EMS mutagenesis in the pea aphid Acyrthosiphon pisum. G3: Genes, Genomes, Genetics, 4, 657–667. https://doi.org/10.1534/g3.113.009639

Tagu D, Hudaverdian S, Le Trionnaire G, Richard G, Gleonnec F, Prunier N, Gauthier JP (2020) Dopamine pathway in the pea aphid. Raw data: doi 10.15454/SHZQBM. url = {https://doi.org/10.15454/SHZQBM}

Tagu D, Hudaverdian S, Le Trionnaire G, Richard G, Gleonnec F, Prunier N, Gauthier JP (2020) Dopamine pathway in the pea aphid.Supplementary data: https://doi.org/10.57745/XHUV1S.

True JR, Edwards KA, Yamamoto D, Carroll SB (1999) Drosophila wing melanin patterns form by vein-dependent elaboration of enzymatic prepatterns. Current Biology, 9, 1382–1391. https://doi.org/10.1016/S0960-9822(00)80083-4

Wang, X, Kang L (2014) Molecular mechanisms of phase change in locusts. Annual Review of Entomology, 59, 225–244. https://doi.org/10.1146/annurev-ento-011613-162019

